# Targeted mutagenesis of Δ5 and Δ6 fatty acyl desaturases induce multiplex-mutagenesis and lipogenesis in Atlantic salmon (*Salmo salar*)

**DOI:** 10.1101/2020.03.12.988873

**Authors:** Yang Jin, Alex K. Datsomor, Rolf E. Olsen, Jon Olav Vik, Jacob S. Torgersen, Rolf B. Edvardsen, Anna Wargelius, Per Winge, Fabian Grammes

## Abstract

With declining wild fish populations, farmed Atlantic salmon (*Salmo salar*) has gained popularity as a source for healthy long-chain highly unsaturated fatty acids (LC-HUFA) including 20:5n-3 and 22:6n-3. However, the introduction of plant-based oil in fish diets has reduced the content of these beneficial LC-HUFA. The capability of biosynthesis of LC-HUFAs depends on fatty acids supplied in diets and the genetic potential residing in the fish. Key proteins involved in LC-HUFA synthesis in salmon include fatty acid desaturases 2 (Fads2). In a recent study we used CRISPR/Cas9 to generate two F0 mutant strains of salmon, 1) *Δ6abc/5*^Mt^ with mutations in *Δ5fads2, Δ6fads2-a, Δ6fads2-b and Δ6fads2-c* genes, and 2) *Δ6bc*^Mt^ with mutations in *Δ6fads2-b and Δ6fads2-c* genes. The CRISPR mutated salmon (crispants) had reduced levels of LC-HUFA and expression of targeted *fads2* genes. In present study we apply whole transcriptome analysis on these *fads2* crispants. Our purpose is to evaluate the genetic mosaicism in *fads2* crispants and the effect these mutations had on other lipid metabolism pathways in fish. Both *Δ6abc/5*^Mt^ and *Δ6bc*^Mt^ crispants demonstrated high percentage of indels within all intended target genes, though different indel types and percentage were observed between individuals. Skipping of a CRISPR-targeted exon was observed in *Δ6fads2-a* gene of *Δ6abc/5*^Mt^ salmon. The *Δ6abc/5*^Mt^ fish also displayed several disruptive indels which resulted in over 100 differentially expressed genes (DEGs) enriched in lipid metabolism pathways in liver. This includes up-regulation of *srebp1* genes as well as genes involved in fatty acid *de-novo* synthesis, fatty acid *β*-oxidation and lipogenesis. Both *elovl5* and *elovl2* genes were not changed, suggesting that the genes were not targeted by Srebp1. The mutation of *Δ6bc*^Mt^ surprisingly resulted in over 3000 DEGs which were enriched in factors encoding genes involved in mRNA regulation and stability.

## 1. Introduction

Atlantic salmon (*Salmo salar* L.) is one of the most beneficial fish species for human consumption since it contains high amounts of long-chain highly unsaturated fatty acids (LC-HUFA) such as docosahexaenoic acid (22:6n-3, DHA), eicosapentaenoic acid (20:5n-3, EPA) and arachidonic acid (20:4n-6, ARA). The high LC-HUFA content in farmed salmon originates mainly from dietary inclusions of marine fish oil and fish meal. However, traditional marine fisheries are exploited to its maximal level and with increasing volume of salmon production, dietary marine oil and meal sources have been gradually diluted over the past decades. Plant oil are used to substitute marine oils in diet, with an increasing levels from 0% of total lipids in 1990 to 19.2% in 2013 [1]. This has resulted in a reduction of LC-PUFA levels in salmon flesh since plant oil do not contain LC-PUFA [2].

Salmon are capable of synthesizing LC-HUFA through elongation and desaturation of α-linolenic (18:3n-3) and linoleic (18:2n-6) acids, and the synthesis is often increased when the fish are given plant oil diet with low LC-HUFA [3]. This explains the fact that salmon can tolerate partially substitution of fish oil with plant oil without negative impact on growth rate, feed conversion or any histopathological lesions [4]. However, the synthesized LC-HUFA in salmon is still not enough to compensate for the reduced LC-HUFA level caused by inclusion of plant oil in diet [2]. This has reduced the nutritional value of salmon for human consumption. One way to solve this is to improve the capacity of LC-HUFA synthesis in salmon to produce higher amounts of LC-HUFA when fed a normal plant oil diet. This however, requires a better understanding of the regulation of genes involved in LC-HUFA synthesis.

The pathways of LC-HUFA synthesis in salmon involves 4 elongases encoded by *elovl2, elovl4, elovl5a* and *elvol5b* and 4 desaturases encoded by *Δ5fads2, Δ6fads2-a, Δ6fads2-b* and *Δ6fads2-c*. All 8 genes have been cloned and functionally characterised through heterologous expression in yeast (*Saccharomyces cerevisiae*) [5, 6]. Both *elovl5a* and *elovl5b* are mainly involved in elongating C18 and C20 fatty acids, while *elovl2* and *elovl4* are involved in elongating C20 and C22 [7, 8, 6]. All four *fads* genes in salmon are homologs to the human *FADS2* gene. In salmon they have separate functions where double bonds are introduced at C5 (*Δ5fads2*) or C6 (*Δ6fads2-a, Δ6fads2-b* and *Δ6fads2-c*) from the carboxyl end [8, 9]. Feeding of plant oil often leads to up-regulation of both *elovl* and *fads2* genes in salmon, which is likely due to the low LC-HUFA content in the diet [10–13].

In addition to the LC-HUFA synthesis genes, many other genes involved in fatty acid *de-novo* synthesis, fatty acid oxidation and cholesterol biosynthesis are also differentially expressed after feeding plant oil [10–13]. It is difficult to conclude the reason for the differential expression of lipid metabolism genes since plant oils are devoid of cholesterol and LC-HUFA and contains high amount of C_18_ PUFA precursors and phytosterols compared to fish oil [14–16]. In a recent study we disrupted LC-HUFA synthesis pathway in salmon by mutating *elovl2* gene using CRISPR/Cas9 technology [17]. In addition to the decreased DHA content in mutant fish, we have identified up-regulation of *fads2* genes as well as several genes involved in fatty acid biosynthesis and lipogenesis [17]. This suggests a systemic change of lipid metabolism regulation in response to the disruption of LC-HUFA synthesis in salmon.

CRISPR/Cas9 technology has recently been used in salmon to edit genes and generate mutants [17–20]. Both guide RNA (gRNA) and Cas9 mRNA are injected to one-cell stage salmon embryos to induce a targeted double-strand break, followed by non-homologous end joining (NHEJ) which generates random insertions and deletions (indels) at the target sites that can leads to a truncated protein. In salmonids, the application. The studies of CRISPR/Cas9 edited salmon have so far been done on the F0 generation because the fish has a long maturation period of 2-4 years. As a mosaic pattern of mutations forms in F0 crispant salmon, which translates into multiple alleles in the fish [17].

We have recently used CRISPR/Cas9 to mutate *fads2* genes in salmon which resulted in down-regulation of targeted genes and lower DHA and EPA contents in tissues [21]. In present study we aimed to further characterize transcriptional regulation of lipid metabolism in *fads2-mutated* salmon by comparing its transcriptomes to wildtype fish. Our study also seeks to provide detailed insights on the effect and distribution of genetic mosaicism in salmon individuals after mutation of *fads2* genes.

## 2. Methods

### 2.1 Generation of CRISPR/Cas9-mediated mutated salmon and feeding experiment

The generation of CRISPR/Cas9-mediated mutated salmon eggs and the corresponding feeding trial was previously published in [21]. In brief, two types of *fads2* mutants were generated with CRISPR/Cas9. Both times a single CRISPR guide RNA (gRNA) was used to target different combinations of *fads2* genes simultaneously: A *Δ6abc/5*-mutated (*Δ6abc/5*^Mt^) salmon strain was generated using a gRNA targeting *Δ6fads2-a* (NCBI Gene ID 100136441), *Δ6fads2-b* (100329172), *Δ6fads2-c* (106584797) and *Δ5fads2* (100136383). A *Δ6bc*-mutated (*Δ6bc*^Mt^) salmon strain was generated targeting *Δ6fads2-b* and *Δ6fads2-c*. Both strains were co-injected with a gRNA targeting the *slc45a2* gene, involved in melanin synthesis [18]. Target sequences of gRNAs were published in Datsomor *et. al*, 2019.

The feeding trial was performed on Atlantic salmon parr of approximate 85 ± 25 g for *Δ6abc/5*^Mt^ salmon, 104 ± 25 g for *Δ6bc*^Mt^ salmon, and 176 ± 34 g for wildtype controls (WT) at the Institute of Marine Research (Matre, Norway). Fish were initially fed a standard commercial diet until start of the experiment. A total of six experimental tanks were used with a common-garden approach, each containing 18 fish consisting of 6 Pit-tagged fish of the *Δ6abc/5*^Mt^, *Δ6bc*^Mt^ and WT. Three tanks were then fed a plant oil diet containing 5% LC-HUFA of total fatty acids in diet, while the remaining three tanks were fed a fish oil diet with 20% LC-HUFA. The fatty acids composition of the diets was shown in detail in [21]. After 54 days of feeding, fish under plant oil diet reached 203 ± 51g for *Δ6abc/5*^Mt^ salmon, 281 ± 52g for *Δ6bc*^Mt^ salmon and 250 ± 62 for WT, while the fish under fish oil diet reached 171 ± 36g, 191 ± 69g and 241 ± 47g for the three groups respectively. Liver and muscle tissues from 6 fish per treatment were then sampled and tissues were flash frozen on dry ice and subsequently stored at −80 °C.

### 2.2 AmpliSeq

To confirm CRISPR/Cas9-induced mutations, AmpliSeq was conducted according to the Illumina protocol (16S Metagenomic Sequencing Library Preparation # 15044223 Rev. B). DNA was isolated from selected individuals from both liver and muscle using DNeasy blood and tissue kits (Qiagen, Hilden, Germany). Primers were designed to specifically amplify the regions around the CRISPR gRNA target sites (Table 1). For each sample the amplicons were generated in singleplex reactions, pooled and then purified using AMPure beads before running index-PCR using the Nextera XT Index Kit (Illumina, San Diego, CA, USA). AmpliSeq libraries were subsequently normalized before sequencing the libraries as 300bp paired-end reads on Illumina MiSeq (Illumina, San Diego, CA, USA) at Centre of Integrative Genetics (CIGENE, Ås, Norway). Raw .fastq reads were quality trimmed using *cutdapt* [22] before aligning them to the salmon genome ICSASG_v2 (https://www.ncbi.nlm.nih.gov/assembly/GCF_000233375.1/) using *bwa mem* [23] and saving files in .bam format. For each sample the proportion of indels for each base in a 25bp window around the target sites was determined using the python3 coverage. py (https://gitlab.com/fabian.grammes/crispr-indel). Additionally we predicted the effect of each indel on the main transcript/protein using *SnpEff* [24].

**Table 1:**
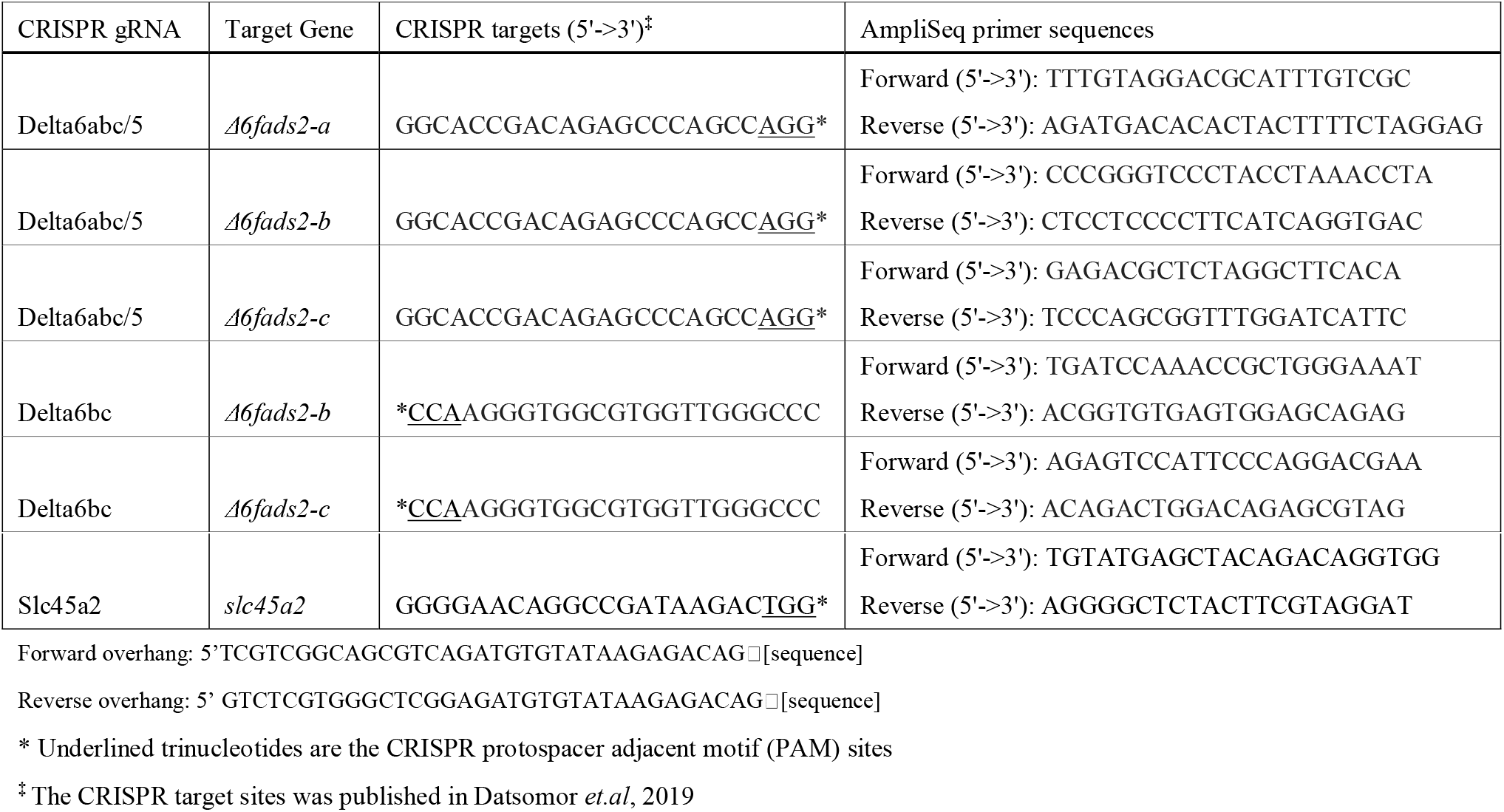
CRISPR gRNA target sequences and AmpliSeq primer sequences.

### 2.3 RNA extraction and library preparation

Total RNA was extracted from liver of individual fish by using RNeasy Plus Universal Mini kit (Qiagen), according to manufacturer’s instruction. RNA concentration and quality were assessed by Nanodrop 8000 (Thermo Scientific, Wilmington, USA) and Agilent 2100 Bioanalyzer (Agilent Technologies, Santa Clara, CA, USA). All samples had RIN values >8.5. RNA-seq libraries were prepared using TruSeq Stranded mRNA Library Prep Kit (Illumina). The libraries were subsequently sequenced using 100bp single-end high-throughput mRNA sequencing (RNA-seq) on an Illumina Hiseq 2500 (Illumina) at Norwegian Sequencing Centre (Oslo, Norway).

### 2.4 Data analysis and statistics

Read sequences were processed using the *bcbio-nextgen* pipeline (https://github.com/bcbio/bcbio-nextgen). In brief reads were aligned to the salmon genome (ICSASG_v2) using *STAR* [25]. The resulting .bam files were subsequently used to generate i) raw gene counts using *featureCounts* (v1.4.4) [26] using the NCBI salmon genome annotation (available for download at http://salmobase.org/Downloads/Salmo_salar-annotation.gff3). ii) exon counts using *DEXSeq* (dexseq_count.py) [27]. In addition reads were mapped directly to the transcriptome using Salmon (v0.10.2) [28]. Raw fastq files and raw gene counts table are publicly available under the accession: E-MTAB-8319 at the ArrayExpress Archive (https://www.ebi.ac.uk/arrayexpress/).

Expression analysis of the genes was performed using R (v3.4.1). Only genes with a minimum counts level of at least 1 count per million (CPM) in 75% of the samples were kept for further differential expression analysis (DEA). DEA was performed between samples from two factors (genotype and diet), using generalized linear model (GLM) method in R package edgeR [29]. The present study focus on three contrasts, *Δ6abc/5*-mutated salmon *versus* WT fed plant oil diet, *Δ6abc/5*-mutated salmon *versus* WT fed fish oil diet, and WT salmon fed plant oil *versus* fish oil diet. Genes with a false discovery rate (FDR), an adjusted *p* value (*q*) <0.05 and absolute log2 fold change (|Log2FC|) >0.5 were considered to be differentially expressed genes (DEGs) between the two test conditions. Subsequently, a KEGG ontology enrichment analysis (KOEA) was conducted using same package edgeR. Hypergeometric test was applied based on number of DEGs compared to total genes annotated to each KEGG pathway, and differences were considered significant when *p* <0.005. The R code which used to generate DEA and KOEA was publicly available at Fairdomhub (https://fairdomhub.org/investigations/242). All figures were made by using R package ggplot2 [30].

## 3. Result and discussion

### 3.1 CRISPR/Cas9 induced mutations

The two strains of Atlantic salmon carrying CRISPR/Cas9-mediated mutations were generated as described earlier [21]. In both strains CRISPR/Cas9 mediated mutations were induced using a single CRISPR gRNA targeting multiple genes (Figure 1A). The gRNA of *Δ6abc/5*^Mt^ salmon targeted *Δ6fads2-a, Δ6fads2-b, Δ6fads2-c* and *Δ5fads2* genes, while the gRNA of *Δ6bc*^Mt^ targeted *Δ6fad2s-b* and *Δ6fads2-c*. Both *Δ6abc/5* and *Δ6bc* mutant salmon were co-injected with a CRISPR gRNA targeting *slc45a2* which induces an albino phenotype and served as visual control in our experiment.

**Figure 1.**
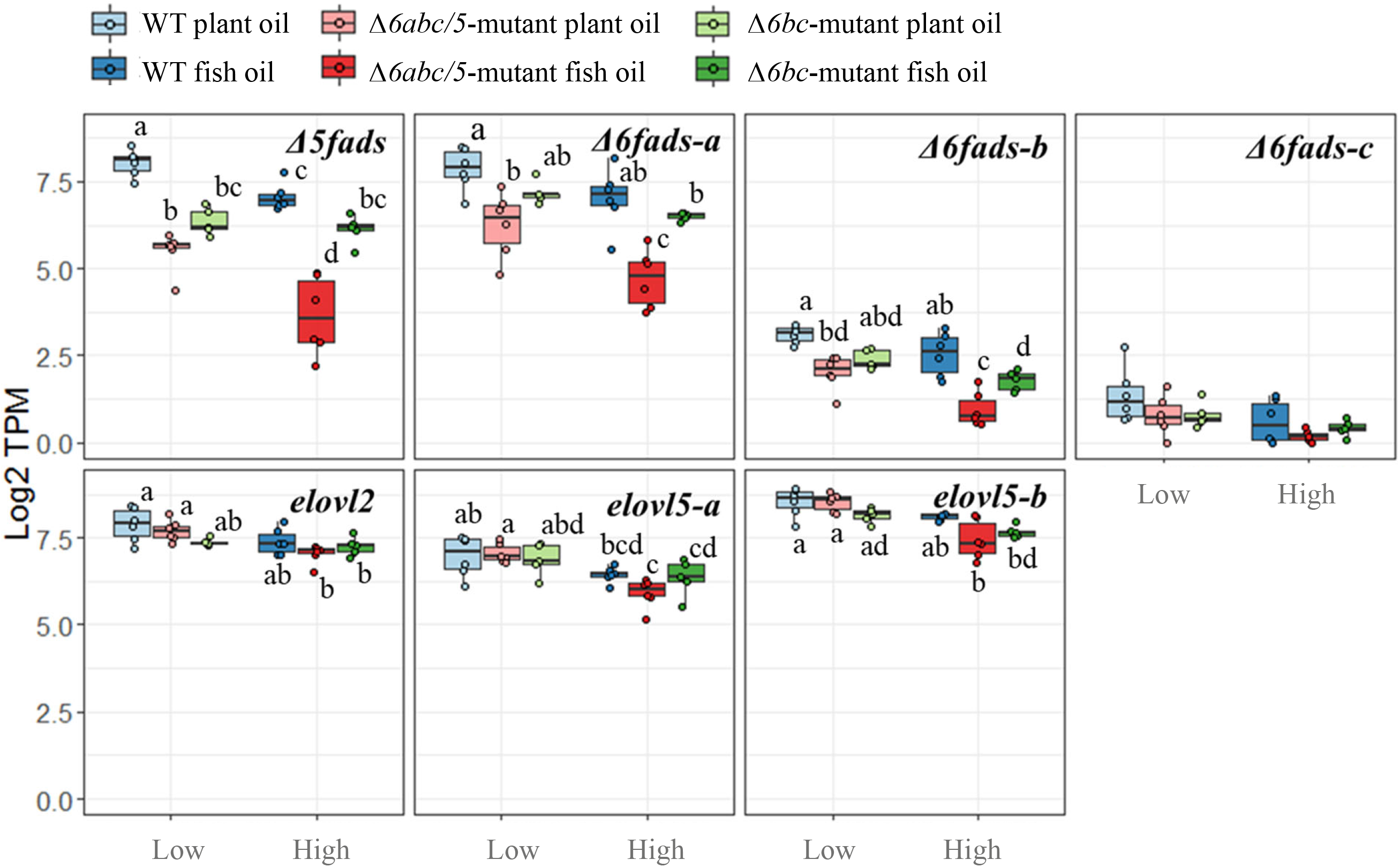
**A**: Circos plot showing the different target sites of the CRISPR gRNAs. Gene Δ*5fads2*, Δ*6fads2-a* and Δ*6fads2-c* have multiple transcripts while yellow boxes indicate exons of each transcript. **B**: Boxplot showing the maximum proportion of insertions/deletions (indels) within the CRISPR gRNA target site as identified by AmpliSeq. Different color indicates liver (L) or white muscle (WM) tissues from WT, Δ*6abc 5* mutant or Δ*6bc* mutant salmon. Each dot indicates L or WM tissue of an individual fish. **C**: Bar plots showing the (SnpEff) predicted impact of the indel on the respective main transcript by individual. Impacts are classified as: HIGH=The variant is assumed to have high (disruptive) impact in the protein; MODERATE=A non-disruptive variant that might change protein effectiveness; LOW=The variant is assumed to be mostly harmless; WT=Wild type/no indel. Each bar of the figure represents data of an individual fish.

CRISPR/Cas9-induced structural mutations at the *fads2* as well as the *slc45a2* genes of fish from both *Δ6abc/5*^Mt^ and *Δ6bc*^Mt^ strains were confirmed by using AmpliSeq. All fish injected with CRISPR/Cas9 carried structural variants at the respective gRNA target sites (Figure 1 B). For all individuals from both CRISPR strains we observed a high degree of mosaicism at each of the respective gRNA target sites (Figure 1B). This suggests that Cas9-induced editing continues after one-cell stage of the embryos. In order to better understand the consequences of the different structural variants on a phenotypic level, we predicted variant effects using SnpEff and summarised the results according to the impact category (Figure 1C). The majority of structural variants across all individuals were predicted to have “high” impact, meaning to have a likely disruptive effect on the protein function. Nevertheless, our analysis also showed that many of the individuals from the two CRISPR strains still carried a considerable amount of the WT genotype (non-CRISPR mutated). Therefore, we believe it is more correct to consider the two resulting CRISPR strains as *fads2* knock-downs rather than knockouts. The *Δ6abc/5*^Mt^ gRNA targeted sequence right after the cytochrome b5-like domain of *fads2* genes, while *Δ6bc*^Mt^ gRNA targeted sequences on exon 1 before all protein domains. Therefore, the out-of-frame mutations in *Δ6abc/5*^Mt^ and *Δ6bc*^Mt^ were expected to disrupt characteristic domains identified in fatty acyl desaturases, though our CRISPR-target sites did not specifically fall within protein domains. These out-of-frame mutations identified by Ampliseq could explain the nonsense-mediated decay (NMD) of the mutant mRNA and impaired biosynthesis of LC-PUFA in *Δ6abc/5*^Mt^ fish [21].

### 3.2 CRISPR/Cas9-induced indels cause *Δ6fads2-a* exon skipping events

Interestingly, we found that CRISPR/Cas9 induced mutations of *Δ6abc/5*^Mt^ gRNA in the *Δ6fads2-a* gene were affecting splicing of exonic part 6 (harbouring the CRISPR target site; exonic part 6 corresponds to exon 4 in transcript: <u>XM_014170212.1</u>; exon 3 in <u>XM_014170213.1</u>). Analysis of exonic-part 6 retention in *Δ6abc/5*-mutated salmon using RNA-seq data revealed mis-splicing of the *Δ6fads2-a* transcript resulting in the skipping of exonic part 6 (Figure 2). Exon skipping caused by CRISPR/Cas9-generated mutations was observed previously in both cell lines [31, 32] and genetically modified organisms including zebrafish [33] and salmon [17]. CRISPR induced mis-splicing is mostly caused by one of two mechanisms: i) indels generated by CRISPR-mutation affects the exon-intron boundaries or ii) indels promote exon skipping by disrupting an exon splicing enhancer or introducing an exon splicing silencer within the targeted exon [34]. However, neither mechanism fits to our study. This was because other *Δ6abc/5*^Mt^ gRNA target sites on *Δ5fads2, Δ6fads2-b* and *Δ6fads2-c* gene contained identical sequences and showed the same distance to exon-intron boundary but did not affect splicing. Nonetheless, the skipping of exon 6 in *Δ6fads2-a* transcripts will result in the production of truncated proteins that lack 37 amino acids, which suggests deleterious effects on protein structure and functions.

**Figure 2.**
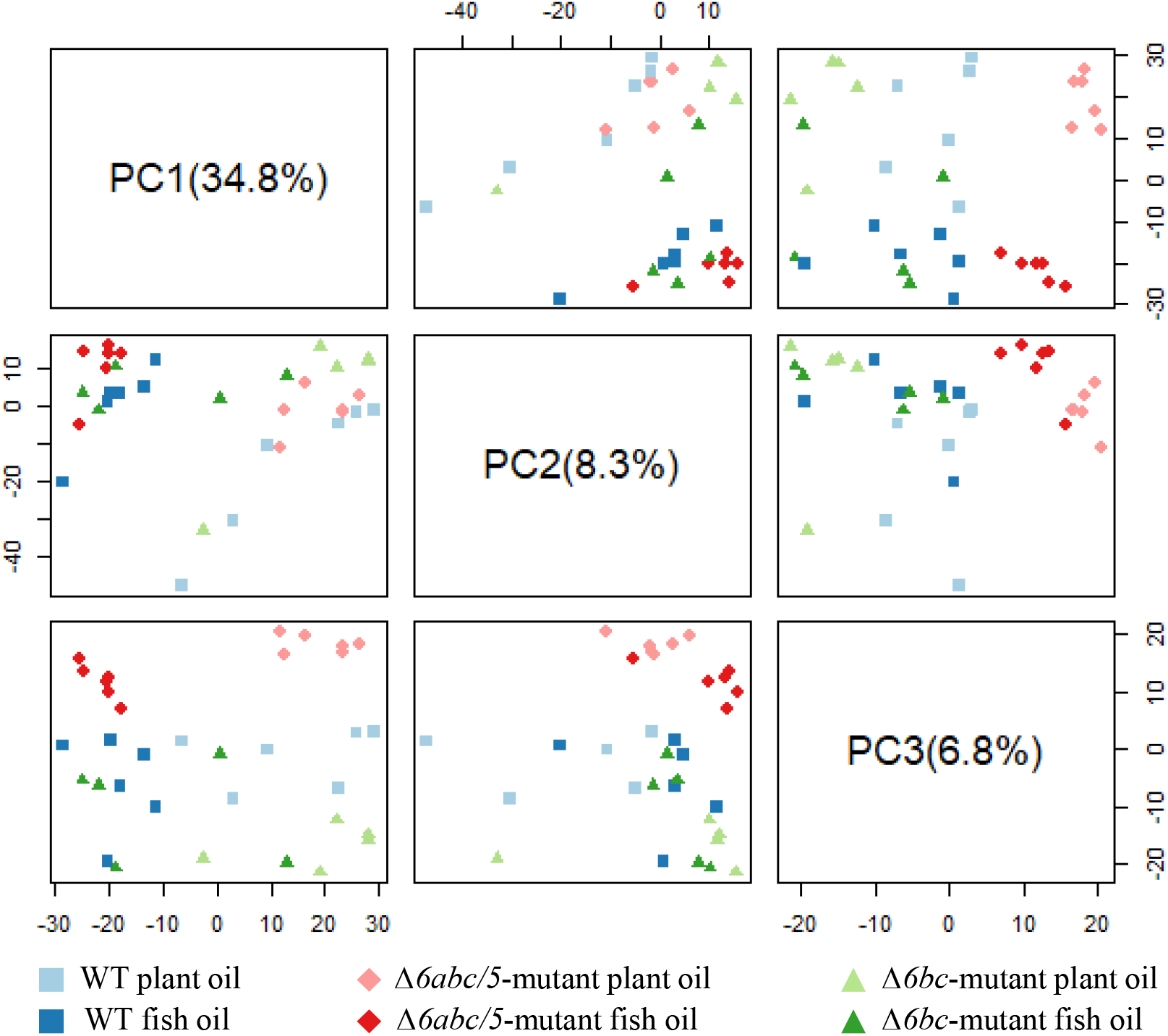
Detection of exon skipping in *Δ6fads2-a* in relation to CRISPR. **A**: Exon structure for the three transcripts encoded by *Δ6fads2-a*. The targeting site (s1) for the *Δ6abc/5^Mt^* gRNA is enlarged and highlighted in red. **B**: Schematic drawing on how aligned RNA-seq reads were used to calculate the percentage of exon retention (PER) for a sample. **C**: Exon skipping was confirmed by using the aligned RNA-seq reads to calculate the PER for each sample (represented as point).

### 3.3 CRISPR-targeted *fads2* genes are down-regulated in *Δ6abc/5* but not in *Δ6bc* salmon

Many of the CRISPR induced structural variants introduce premature termination codons likely to trigger mRNA degradation by nonsense-mediated decay (NMD) [35]. Indeed, we found that CRISPR-targeted *Δ5fads2, Δ6fads2-a* and *Δ6fads2-b* genes were strongly down-regulated (*q*<0.05) in *Δ6abc/5*^Mt^ salmon compared to WT regardless of the dietary treatment (Figure 3). In *Δ6bc*^Mt^ salmon, the CRISPR-targeted *Δ6fads2-b* gene was down-regulated compared to WT, but the levels of down-regulation were less clear than in *Δ6abc/5*^Mt^ salmon. Surprisingly, the expression *Δ5fads2* and *Δ6fads2-a* genes was also down-regulated in *Δ6bc*^Mt^ salmon, though both genes were not targeted by *Δ6bc*^Mt^ gRNAs. The expression of *Δ6fads2-c* gene was generally very low and was not likely play a major role in salmon liver. This low level expression may also explain that *Δ6fads2-c* was not affected by CRISPR mutation (Figure 3). The expression of other genes in LC-HUFA synthesis pathways, *elovl2, elovl5-a* and *elovl5-b* genes was stable between *Δ6abc/5*^Mt^, *Δ6bc*^Mt^ and WT salmon.

**Figure 3.**
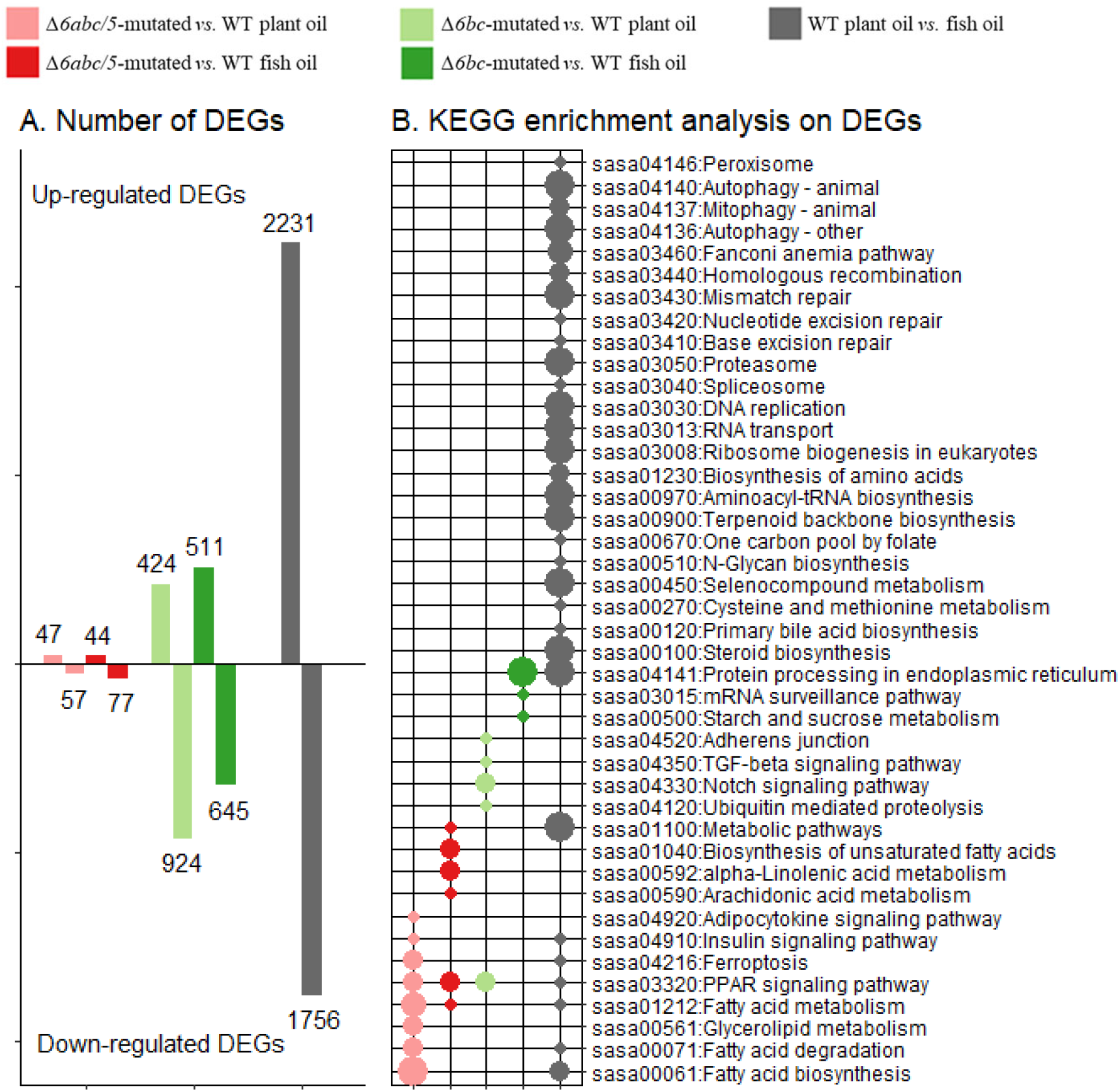
Expression of LC-HUFA synthesis genes in wildtype (WT), *Δ6abc/5^Mt^* and *Δ6bc^Mt^* salmon fed with either plant oil or fish oil diet. Gene expression are shown in transcript per million (TPM) value which is raw counts normalised by both library size and mRNA length. Different letter indicates genes which were differentially expressed (*q*<0.05 & |log2FC|>0.5).

The NMD-mediated mRNA degradation, absence of exon 6 in *Δ6fads2-a* transcripts, and other CRISPR-induced mutations such as out-of-frame mutations are expected to produce non-functional enzyme proteins that would ultimately disrupt LC-HUFA biosynthesis in the fish. Indeed, analysis of tissue composition of LC-HUFA coupled with assay of desaturation and elongation activities in liver showed clear impacts of the CRISPR-mutations. The mutation of *Δ6abc/5* genes in salmon resulted in significant reduction of DHA and EPA compared to WT [21]. On the other hand, we observed effects of background wildtype alleles in the *Δ6abc/5*^Mt^ salmon (Figure 1B and C) accounting for limited but measurable desaturation activities [21].

Low levels of LC-HUFA levels often induce hepatic expression of *Δ5fads2* and *Δ6fads2-a* genes as was shown in our previous *elovl2*-mutated salmon [17]. On the other hand, reduced DHA level has little effect on the expression of *elovl5* and *elovl2* genes as was shown in present *Δ6abc/5*^Mt^ salmon (Figure 3). Higher expression of *elovl2* and *elovl5* genes that often observed in fish fed plant oil compared to fish oil diet (Figure 3) [36, 37], which was more likely caused by other differences between the two diets, such as cholesterol levels.

### 3.3 Transcriptional changes in liver after mutating *fads2* genes

An average of 29 million reads were mapped on to the salmon genome ICSASG_v2. From a total of 55304 annotated genes, 23114 genes had at least 1 count per million (CPM) in 25% of the samples, and were considered for subsequent analysis. By applying principle component analysis (PCA) on Log2 CPM of the top 1000 most variant genes, we identified a clear separation of plant oil and fish oil samples between PC1 (explaining 34.8% of the observed variation) and PC2 (8.3%) as well as a separation of WT and *Δ6abc/5*^Mt^ samples between PC2 and PC3 (6.8%) (Figure 4). Although not as strong we also found a clear tendency for separation of WT and *Δ6bc*^Mt^ samples between PC2 and PC3. Plant oil diet and CRISPR-mutation seemed to have different impact on gene transcription in salmon liver, though both the diet and mutation have generated low levels of LC-HUFA in the fish body.

**Figure 4.**
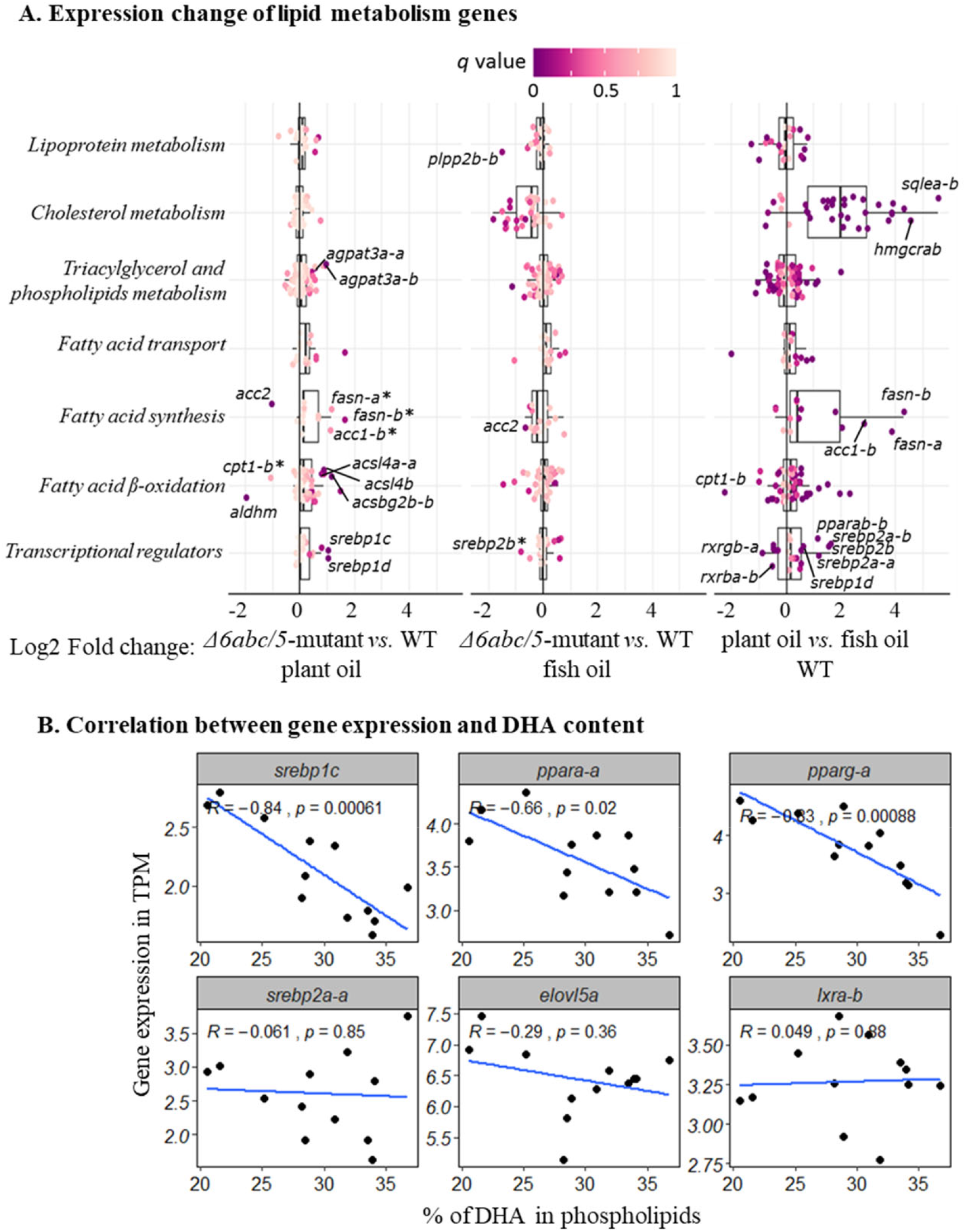
Principle component analysis (PCA) on Log2 count per million (CPM) of the top 1000 most variant genes between all liver samples. Different colors represents genetic groups of WT, *Δ6abc/5-mutated* and *Δ6bc*-mutated salmon, while the color intensity represents different dietary treatments of either plant oil (low HUFA) diet or fish oil diet (high HUFA).

Differential expression analysis (DEA) was done by contrasting crispat and WT salmon separately under plant oil and fish oil diets. This resulted in 121 differentially expressed genes (DEGs, *q*<0.05 & |log2FC| >0.5) in *Δ6abc/5*^Mt^ salmon compared to WT when fed fish oil diet, while 104 DEGs were found between crispat and WT salmon under plant oil diet (Figure 5 A). Surprisingly, more DEGs were found in *Δ6bc*^Mt^ salmon compared to WT. This includes 1665 genes identified in crispat salmon when fed fish oil diet and 2041 DEGs identified in salmon fed plant oil diet. A total number of 3863 DEGs was found in WT salmon fed plant oil diet compared to fish oil.

**Figure 5.**
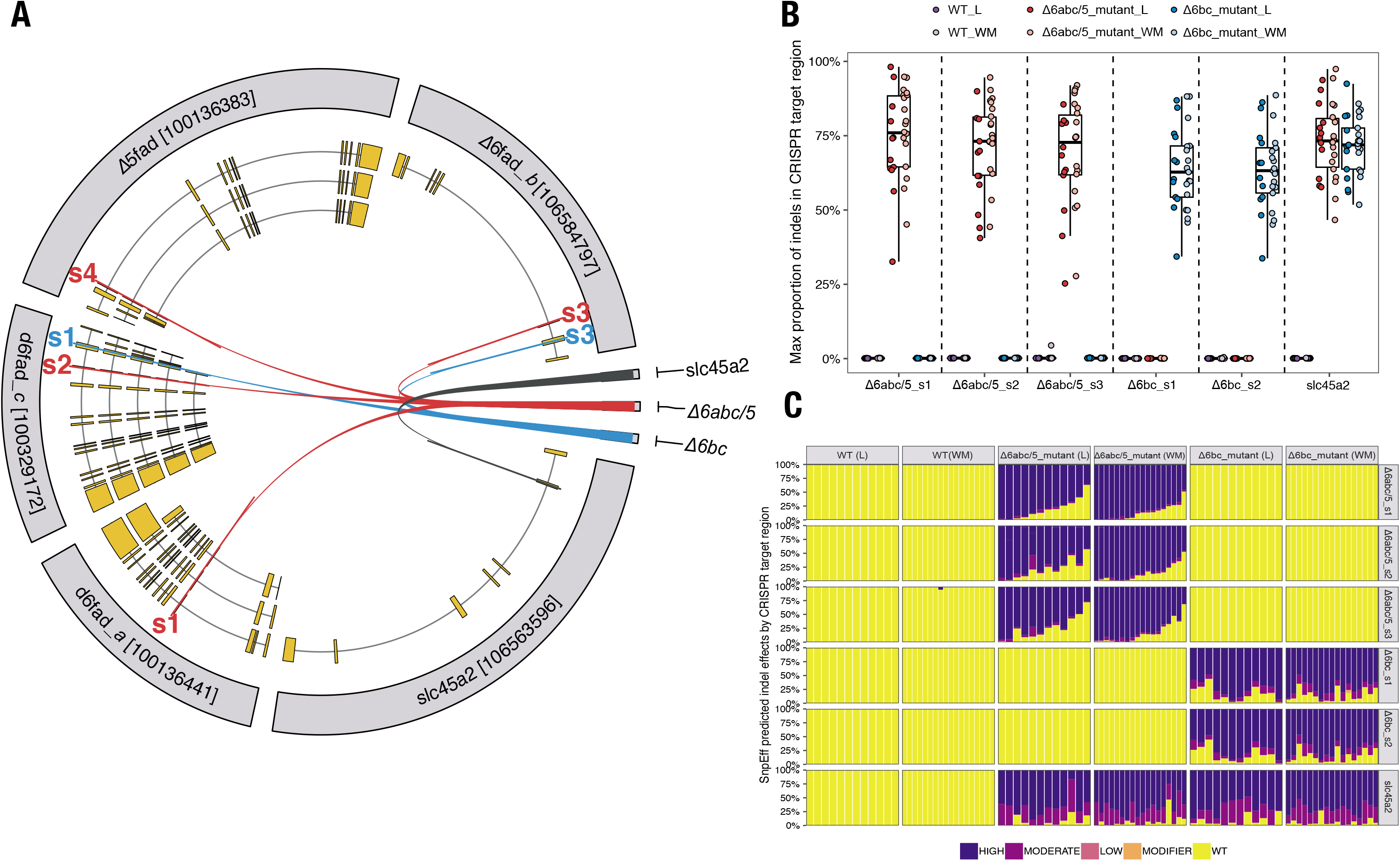
Differential expression analysis in liver between wildtype (WT) and mutated salmon. **A)** Number of up-regulated and down-regulated differential expressed genes (DEGs, q<0.05 & |log2FC| >0.5) either between WT and *Δ6abc/5*-mutated salmon, or between WT and *Δ6bc*-mutated salmon, or between WT salmon fed plant oil and fish oil. **B)** Significantly (*p*<0.005) enriched KEGG pathways of the DEGs. Hypergeometric test was applied based on the number of DEGs versus total genes annotated to each KEGG pathway.

To further understand the functions of DEGs between crispat and WT salmon, we conducted a KEGG enrichment analysis by comparing the number of DEGs to the total number of genes in each KEGG pathway (Figure 5 B). The DEGs of *Δ6abc/5*^Mt^ salmon were not only enriched in fatty acid metabolism pathway, but also peroxisome proliferator-activated receptors (PPAR) signalling pathway which is involved in many metabolic pathways including fatty acid synthesis and catabolism [38]. This suggests PPAR to be key transcription regulators for fatty acid metabolism in salmon. Their differential regulation was likely caused by the decreased EPA and DHA and consequential accumulation of 18:3n-3 and 18:2n-6 after disruption of LC-HUFA synthesis pathway [21]. Accumulated 18:3n-3 and 18:2n-6 could not be synthesised further to DHA and EPA after disruption of *fads2* genes, but were alternatively consumed in *β*-oxidation which was mostly likely activated by PPAR transcription factor [38]. Similar enrichment of fatty acids metabolism and PPAR signalling pathways was also found in the DEGs between WT salmon fed plant oil and fish oil (Figure 5 B). Additionally, sterol biosynthesis pathway was enriched for DEGs between WT salmon fed plant oil and fish oil but was not enriched for the DEGs between *fads2* mutants *versus* WT fish (Figure 5 B). Indicating that the LC-HUFA level and PPAR has little effect on cholesterol biosynthesis in salmon, which should be more likely regulated by other biochemical signals such as low cholesterol level and other transcription factors including sterol regulatory binding protein 2 (SREBP2) [11, 12, 14]. More detailed discussion on PPAR and SREBP regulations will be shown later in this paper. Many other pathways were also enriched for the DEGs of WT fed plant oil *versus* fish oil, such as amino acid biosynthesis and RNA transport. This suggests that dietary inclusion of plant oil has more complex impact on salmon than just reducing LC-HUFA and cholesterol levels in the fish body. Our study has successfully separated the effect of low LC-HUFA level from other effects of plant oil inclusion, however more research is required to understand the complete regulatory network in response to the change of plant oil in diet. Surprisingly, no lipid metabolism pathways were enriched in *Δ6bc*^Mt^ salmon compared to WT, regardless of dietary LC-HUFA level. This was in accordance to the fatty acid composition in liver, where no significant difference was found between *Δ6bc*^Mt^ salmon and WT [21]. The DEGs were likely more enriched in mRNA regulation pathways, including mRNA surveillance and spliceosome pathways. One possible explanation could be the mutation of ncRNA which has function on transcription regulation or immune system [39]. Nevertheless, the reason for the high number of DEGs in *Δ6bc*^Mt^ salmon and their enriched pathways needs to be further investigated.

### 3.4 Expression of lipid metabolism genes in response to *Δ6abc/5* mutation

Due to many unexpected and lipid metabolism unrelated DEGs found in *Δ6bc*^Mt^ salmon, only *Δ6abc/5*^Mt^ fish were included for further transcriptomic analysis to understand the transcriptional regulation of lipid metabolism after disrupting LC-HUFA synthesis genes. Here we discussed DEGs of lipid metabolism pathways that enriched in *Δ6abc/5*^Mt^ *versus* WT salmon, aiming to understand the regulatory network of lipid metabolism genes in response to *Δ6abc/5*^Mt^. The *Δ6abc/5* mutant showed 14 (13.4%) differentially expressed lipid metabolism genes when fed plant oil diet, while less (7 genes, 5.8%) lipid DEGs were identified in salmon fed fish oil diet (Supplementary Table 1). The higher numbers of DEGs in *Δ6abc/5*^Mt^ salmon fed the plant oil diet suggest a compensatory response to the combined effects of impaired endogenous LC-HUFA biosynthesis and reduced dietary LC-HUFA levels. On the other hand, the reduced number of lipid DEGs in *Δ6abc/5*^Mt^ salmon fed fish oil diet suggests an impact of dietary LC-HUFA levels on gene transcription, most likely an end-product-mediated inhibition. Nevertheless, 4 lipid DEGs were identified in *Δ6abc/5*^Mt^ fish fed both plant oil and fish oil experimental diets including *Δ5fad, Δ6fad-a, abcd1* and *acc2*. Besides the two CRISPR-targeted genes, the down-regulation of *acc2* and up-regulation of *abcd1* suggests an increase of fatty acid *β*-oxidation pathway for energy expenditure after CRISPR-mutation [40].

Sterol regulatory element binding proteins (SREBPs) are suggested to be involved in regulating lipid metabolism in both mammals and fish [41, 42]. Atlantic salmon has four *srebp1* paralogous genes, *srebp1a, srebp1b, srebp1c* and *srebp1d* which are all orthologs of the zebrafish *srebp1* gene (Supplementary Table 1). Both *Δ6abc/5*^Mt^ and low LC-HUFA diet resulted in increased transcription of all four *srebp1* genes in salmon (Figure 6 and Supplementary Table 1). The transcription of the *srebp1* genes was negatively (*p* <0.05) correlated to the DHA level in phospholipid. On the other hand, transcription of *srebp2* genes were not up-regulated in mutated *versus* WT salmon, and are not correlated to DHA level (Figure 6 B). The different regulation of *srebp1* and *srebp2* transcription is consistent with previous studies on mammals, suggesting that *srebp1* transcription is regulated by DHA levels in salmon, while *srebp2* transcription is more likely to be induced by low cholesterol level in the plant oil diet [41].

**Figure 6.**
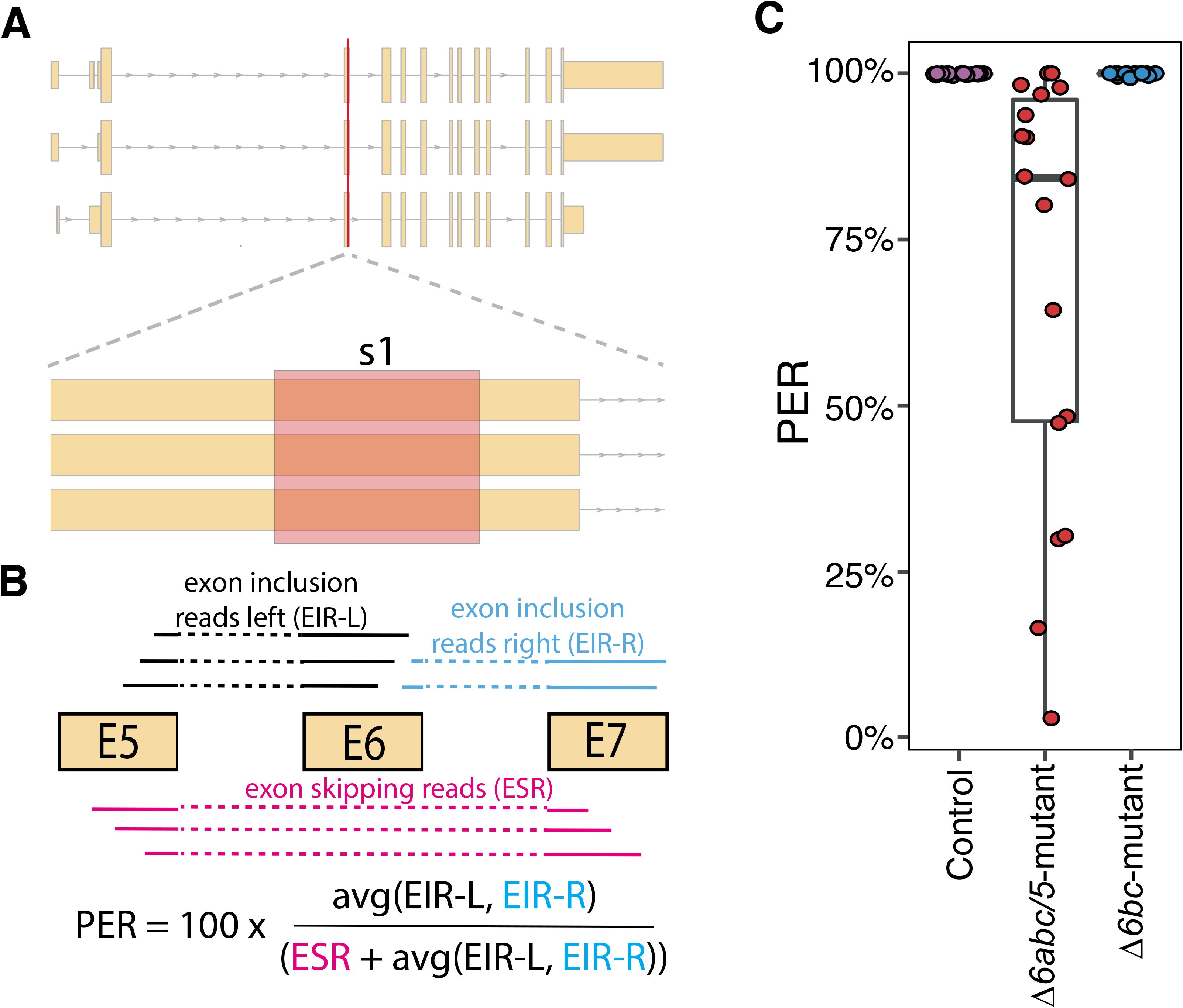
Expression change of liver genes involved in lipid metabolism after CRISPR mutation. **A)** Expression changes of genes in Log2 fold change between mutated and wildtype salmon. Differentially expressed genes (DEGs, q<0.05 & |log2FC|>0.5) are labelled, except three genes with asterix (*) which had high log2 fold change but not significant (*q*>0.05) **B)** Correlation between gene expression and DHA content in phospholipid. Data of DHA measurement was aquired from Datsomor *et.al*, 2019.

By comparing salmon gene promoter sequences to 6 transcription factor binding sites databases (CISBP, HUMAN.H10MO.B, HT-SELEX2, HumanTF, JASPAR, TRANSFAC), we identified 235 lipid metabolism genes, with potentiall sterol regulatory elements (SRE), the Srebp binding sites, between −1000bp to 200bp from transcription starting sites (Supplementary Table 2). This includes *Δ5fad, Δ6fad-a, elovl5a, elovl5b* and *elovl2* which are the major genes in LC-HUFA synthesis pathway. Recent study showed that CRISPR/Cas9-mediated editing of *elovl2* in salmon has increased transcription of *srebp1, Δ6fad* and *Δ5fad* genes together with decreased LC-HUFA content, supporting the regualtion of *fads2* genes by Srebp-1 transcription regulator [17]. However, salmon Srebp-1 is unlikely to induce *elovl5* and *elovl2* since their expression were stable while *srebp1* expression was up-regulated in *Δ6abc/5*^Mt^ compared to WT salmon fed plant oil. The *elovl5* genes were also stable in *elovl2*-mutated salmon [17]. One possible reason is that the SRE in promoter regions of *elovl5* and *elovl2* genes may be more efficient for binding Srebp-2 rather than Srebp-1 [43], or other that transcription factors such as liver X receptor (LXR) are responsible stimulation of *elovl* genes in salmon under plant oil diet. On the other hand, mammlian SREBP-1 can target both fatty acid desaturase (*FADS2*) and elongase (*ELVOL5*) genes and regulate LC-HUFA synthesis [44, 45].

To further investigate the relationship between those key transcription factors and lipid metabolism genes, we compared the expression changes of the 230 lipid metabolism genes except LC-HUFA synthesis genes, either between mutated and WT salmon fed plant oil, or between mutated and WT salmon fed fish oil, or between WT salmon fed plant oil and fish oil (Figure 6A). Several *agpat3* and *acsbg* genes were significantly (*q*<0.05 & |log2FC| >0.5) up-regulated in plant oil mutated salmon together with up-regulated *srebp1*. The function of Srebp-1 transcription factor in salmon is likely similar to its function in mammals, which works as a key transcription factor for hepatic lipogenesis, and *agpat3* and *acsbg* genes are likely the key target genes of salmon Srebp-1. Same *acsbg, agpat3* and *srebp1* genes were also up-regulated when the *elovl2* gene was CRISPR-mutated in salmon, confirming an increase of fatty acid acylation and lipogenesis in response to decreased tissue DHA content [17]. Other typical mammlian SREBP-1 targets, *fasn, acc1* and *elovl6* genes of fatty acid synthesis and elogation pathways were also up-regualted, but not significant (*q*>0.05) in mutated salmon compared to WT under plant oil diet (Figure 6). However, the transcriptional increase of these genes were much higher and significant (*q*<0.05) in WT salmon fed plant oil diet compared to fish oil. This means that the genes of fatty acids synthesis and elongation in salmon were not merely targeted by Srebp-1, but by other transcription factors likely Srebp-2 [41] or Ppar-γ [46]. Genes of cholesterol metabolism including *hmgcrab, mvd-a* and *sqlea-a* were only highly up-regualted in WT fed plant oil diet *versus* fish oil, while no transcription change was observed in *Δ6abc/5*^Mt^ *versus* WT salmon. Several studies have found up-regualtion of choleserol biosyntheis and *srebp2* genes in salmon fed plant oils [11, 12, 14]. The present study has supported that the relationship between *srebp2* and cholesterol biosynthesis genes is quite conserved as in salmon as in mammals, and suggests that the SREBP binding sites of cholesterol biosynthesis genes were *srebp2*-specific [41].

CRISPR/Cas9-mediated mutation of *fads2* genes in *Δ6abc/5* also affected the fatty acid *β*-oxidation pathway in salmon. This was indicated by a strong down-regulation of *acc2* gene following *Δ6abc/5*^Mt^ (Figure 5). Unlike the *ACC1* gene which is mostly involved in *de-novo* fatty acid synthesis in cytosol, the *ACC2* gene in mammals produce mitochondria-associated malonyl-CoA which is a negative regulator of CPT1 and inhibits mitochondria *β*-oxidation [47, 48]. Therefore, the down-regualtion of *acc2* in *Δ6abc/5*^Mt^ salmon could suggest an increased fatty acid *β*-oxidation after disrutpion of LC-HUFA sythetic pathway. This could be regulated by PPAR which is key regualtor of fatty acid catabolism [38]. Similar to *srebp1*, we also found negative correlation between DHA level and two *ppara-a* genes, though its expression was not changed after *Δ6abc/5* mutation. As PUFA and their derivatives are known natural ligands of PPAR, the activation of PPAR and their target genes including fatty acid *β*-oxidation may not rely on increased transcirption of PPAR genes [49]. The increased *β*-oxidation was probably due to accumulation of 18:3n-3, 18:2n-6, and other intermediate fatty acids in LC-HUFA synthesis pathway which cannot be synthesised further to DHA and EPA after disruption of *fads2* genes but were alternatively consumed in *β*-oxidation [21]. Feeding of plant oil diet also induced *cpt1a* and *abcd1*, which are key genes involved in import of fatty acid into mitochondria and peroxisome for catabolism. However, a paralog gene *cpt1b* were down-regulated both after *fads2*-mutation and feeding plant oil diet. The reason for the down-regualtion is unclear and whether it would affect fatty acid *β*-oxidation needs to be further investigated. One possible explanation is that malonyl-CoA produced by *acc1* or *acc2* is less organelle-specific in salmon, and that *cpt1b* gene could be inhibited by malonyl-CoA produced by *acc1* in *de-novo* fatty acid synthesis.

## 4. Conclusions

CRISPR-Cas9 can be employed efficiently to mutate the multiple *fads2* genes, simultaneously, in salmon. However, mosaic effects are common, embodied by different indels among tissues and individuals. The exon skipping, found in the *Δ6fads2-a* gene during transcription, was predicted to result in the production of truncated proteins and strengthen the CRISPR-induced disruption of LC-HUFA synthesis in *Δ6abc/5*^Mt^ salmon. Down-regulation of the targeted *Δ5fad, Δ6fad-a* and *Δ6fad-b* genes were found in liver, which likely cause a decrease of LC-HUFA synthesis. On the other hand, the transcription of *elovl5a, elovl5b* and *elovl2* genes in LC-HUFA synthesis pathway was not affected. Since *srebp1* genes were up-regulated in *Δ6abc/5*-mutated salmon the *elovl* genes were not likely regulated by this transcriptional factor. Increased *de-novo* fatty acid synthesis and lipogenesis was observed after *Δ6abc/5*^Mt^ and could also be regulated by SREBP1. In addition, the level of transcriptional changes of *fasn* and *acc1* genes involved in fatty acid synthesis were much higher when the fish was fed plant oil diet as compared to fish oil. This suggests that these genes were regulated by one or more transcriptional factors in addition to SREBP1. PPAR or SREBP2 are likely candidates. Increased fatty acid *β*-oxidation was also observed after *Δ6abc/5*^Mt^ and was likely regulated by PPAR. The CRISPR-mutation of *Δ6bc*^Mt^ genes surprisingly revealed over 3000 DEGs in liver of salmon, and the DEGs were not enriched in any lipid metabolism pathways. The reason for the high number of DEGs in *Δ6bc*^Mt^ salmon was unclear and needs to be further investigated.

## Acknowledgements

The RNA-Seq and data analysis were financed by the Research Council of Norway (DigiSal, grant number 248792). We would like to thank Matilde Mengkrog Holen and Centre for Integrative Genetics (CIGENE) for help during RNA-Seq sample preparation.

## Author contributions

### Conceptualization

Yang Jin, Alex Datsomor, Rolf Edvardsen, Jacob Torgersen, Per Winge, Fabian Grammes, Rolf Erik Olsen

### Data Curation

Yang Jin, Fabian Grammes

### Formal Analysis

Yang Jin, Fabian Grammes

### Funding Acquisition

Rolf Edvardsen, Jacob Torgersen, Per Winge, Jon Olav Vik, Anna Wargelius

### Methodology

Yang Jin, Fabian Grammes

### Resources

Alex Datsomor, Rolf Edvardsen, Anna Wargelius, Per Winge, Fabian Grammes, Rolf Erik Olsen

### Visualization

Yang Jin, Fabian Grammes

### Writing – Original Draft Preparation

Yang Jin, Fabian Grammes

### Writing – Review & Editing

Yang Jin, Jon Olav Vik, Anna Wargelius, Alex Datsomor, Rolf Edvardsen, Jacob Torgersen, Per Winge, Fabian Grammes, Rolf Erik Olsen

